# Optimising the tilt-increment for in situ cryo-electron tomography

**DOI:** 10.1101/2025.08.20.671201

**Authors:** Maarten W. Tuijtel, Tomáš Majtner, Beata Turoňová, Martin Beck

## Abstract

Cryo-electron tomography (cryo-ET) enables high-resolution, three-dimensional imaging of cellular structures in their native, frozen state. However, image quality is limited by a trade-off between angular sampling and radiation damage. Therefore, the choice of the angular increment during data collection is a critical parameter that affects tomogram quality and downstream analyses. Optimising this increment is challenging due to the high demands on microscope time, storage, and computation. In this study, we systematically evaluated tilt increments of 1°, 2°, 3°, 5°, and 10° using lamellae from *Dictyostelium discoideum* cells. We found that at a constant total electron dose, finer tilt increments (1–3°) produced better-aligned tomograms with higher signal-to-noise ratios and improved outcomes in template matching and subtomogram averaging. A 3° increment emerged as the optimal balance between data quality, alignment accuracy, dose per image, and processing efficiency. This practical recommendation supports both high-throughput and high-resolution structural studies and can guide future cryo-ET data acquisition strategies.

## Introduction

Cryo-electron tomography (cryo-ET) (*1*) is a powerful technique to study virions (*2*), thin bacterial cells (*3*), isolated organelles, larger eukaryotic cells (*4–6*), and even tissue (*7*, *8*) with molecular resolution. It captures the 3-dimensional (3D) electrostatic potential of the specimen under scrutiny and enables the structural analysis of macromolecular complexes within their native context. However, it is also limited by a number of technical parameters. One of these is the maximum electron dose that can be applied to biological specimens before irreversible damage occurs, which is about 100-150 e^-^/Å^2^ for most eukaryotic cells(*9*, *10*).

For simplicity, the cryo-ET workflow can be described in terms of an experimental and two subsequent computational steps, which are relevant to this study. (i) Multiple two-dimensional (2D) projection images are acquired that capture the respective biological specimen from different angles. (ii) These individual projections, which are also referred to as tilt images, need to be computationally aligned to correct for spatial displacements that occur due to mechanical imperfections of the microscope stage. (iii) From the resulting aligned tilt series of projection images, a three-dimensional (3D) reconstruction is computed.

The maximum electron dose influences all three of these steps. (i) The dose has to be distributed over all tilt images and therefore the angular increment is a key experimental parameter for the data acquisition. (ii) The dose contained in an individual projection determines how well it can be computationally aligned with its neighbouring images. Both (i) the data acquisition, and (ii) the alignment of images, impact on the quality of (iii) the reconstruction: The smaller the angular increment used during data acquisition, the lower the signal in each individual tilt image, but the finer the angular sampling of the resulting 3D reconstruction. Inversely, the more accurate the alignment of the tilt images, the better the accuracy of the reconstruction. Since both parameters influence the outcome of the cryo-ET workflow in an interdependent way, it is difficult to make predictions about the optimal tilt increment.

These considerations have led to the notion that there is a trade-off between the number of tilts that can be collected and the electron dose that can be allocated per tilt image. This caveat needs to be considered for every cryo-ET data acquisition session, and different tomographic acquisition schemes have been proposed to deal with this phenomenon (*11–15*). Since accumulated radiation dose progressively degrades high-resolution information, this motivated the development of the dose-symmetric tilt scheme, which prioritizes acquisition of low-tilt images early to better preserve high-resolution information(*11*, *16*). However, to the best of our knowledge, this trade-off has never been systematically addressed and empirically validated, possibly due to the extensive experimental and computational effort associated with such investigations. The respective decision regarding acquisition parameters is thus often made on an arbitrary basis.

The alignment is thus a critical step in the cryo-ET workflow, and it is typically performed in two stages. First, coarse alignment is achieved by cross-correlating the entire projection images to correct for large lateral shifts introduced during image acquisition. Subsequently, the images are subdivided into overlapping patches, and local cross-correlations are computed between adjacent projection images to estimate relative shifts. These patch-derived shifts are then refined using a least-squares fitting procedure to determine optimal alignment parameters (*17*).

The challenges during the 3D reconstruction can be explained using the central slice theorem (*18*), which states that the Fourier transform of each 2D projection (tilt image) corresponds to a central slice through the 3D Fourier transform of the final tomogram. This illustrates the sampling challenge that is inherent to cryo-ET. The sample thickness (*D*) (or the resulting thicknesses of all central slices in Fourier space), and the finite number of 2D projections (*<u>n</u>*) that are used for the reconstruction, result in regions between the tilts that also remain incompletely sampled beyond a defined spatial frequency. The Crowther criterion conceptualises this phenomenon and provides a means to calculate the maximum attainable resolution (*d*) based on angular sampling (*19*, *20*), with:

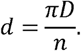

Similarly, the signal-to-noise ratio (SNR) in 3D reconstructions is described by the dose-fractionation theorem (*21*) which states that the statistical significance of each reconstructed voxel depends solely on the total applied electron dose, irrespective of the number of projection images, provided that perfect alignment between the projections is achieved (*22*). Thus, when the total dose is kept constant, the SNR in the tomogram depends on the accuracy of the alignment.

Tomograms are often further data mined by locating specific proteins or protein complexes in the crowded cellular environment, using, for example, template matching (TM) approaches (*23*, *24*), or alternatively, machine-learning based feature recognition (*25*, *26*). For *in situ* structural analysis, the respective particles are extracted, aligned, and averaged, a process called subtomogram averaging (STA) (*27*, *28*). These processing routines are critically important to understand the function of large macromolecular assemblies inside cells (*29*, *30*). Therefore, the outcome of TM and STA in terms of accuracy or resolution, respectively, has been used as a metric to score the technical quality of tomograms and the suitability of data acquisition schemes for specific applications (*31*). During STA processing, the alignment of the projections can be refined on a per-particle basis (*32–37*), which is instrumental for achieving high-resolution structures in the sub-4 Å-range (*5*, *10*, *32*, *38*).

According to the above-discussed notion that there should be a trade-off between angular sampling and radiation damage, we reasoned that there should be an optimal balance between angular sampling and tilt image alignment and, thus, tomogram quality. To address this, we acquired data on lamellae of *Dictyostelium discoideum cells* using various tilt increments, both typical for the field, and beyond. We utilised TM and STA as metrics to determine the optimal tilt increment for *in situ* cryo-ET. Surprisingly, we found that tilt-series alignment improves with smaller angular increments, which, in line with the dose-fractionation theorem, benefits the SNR of the tomograms, despite the decreasing signal in the individual tilt images. Tomographic reconstruction, TM, and STA were successful for all imaging conditions, however, visual tomogram quality, TM accuracy and STA resolution suffered from the use of higher increments. An increment of 3 degrees emerged as the optimal balance between data quality, alignment accuracy, dose per image, and processing efficiency. The results described herein can serve as a guide for the community to choose data acquisition parameters for cryo-ET data collection.

## Results

### A dataset for benchmarking tilt increments in cryo electron tomography

To study the effect of the tilt increment, we collected several cryo-ET datasets using SerialEM (*39*) on lamellae from *Dictyostelium discoideum* cells, varying the tilt increment from 1, 2, 3, 5, to 10 degrees using the dose-symmetric tilt-scheme (*11*) (for details on the sample preparation and microscopy data acquisition, see Methods). During acquisition, the tilt range (from −60 to +60 degrees relative to the lamella plane) and the total dose (120-130 e^-^/Å^2^) were kept constant for each condition. As a consequence, the number of images in a tilt series and the dose per tilt image were varied for each condition (for full details regarding data acquisition parameters, see Table 1). Both data storage requirements and beam time needed per tomogram differed by nearly an order of magnitude (Table 1). After acquisition, tilt series were pre-processed, aligned using patch-tracking, and tomograms were reconstructed in IMOD (*17*) (see Methods). For most datasets, image acquisition was largely complete, with the exception of the 1-degree dataset, which showed a markedly higher proportion of missing images at high tilt angles (SFig. 1). Closer inspection revealed that many of these images were not acquired, as SerialEM applies built-in safeguards (e.g. autofocus inconsistency or insufficient image counts) that can abort a tilt-series branch before completion.

**Fig. 1.**
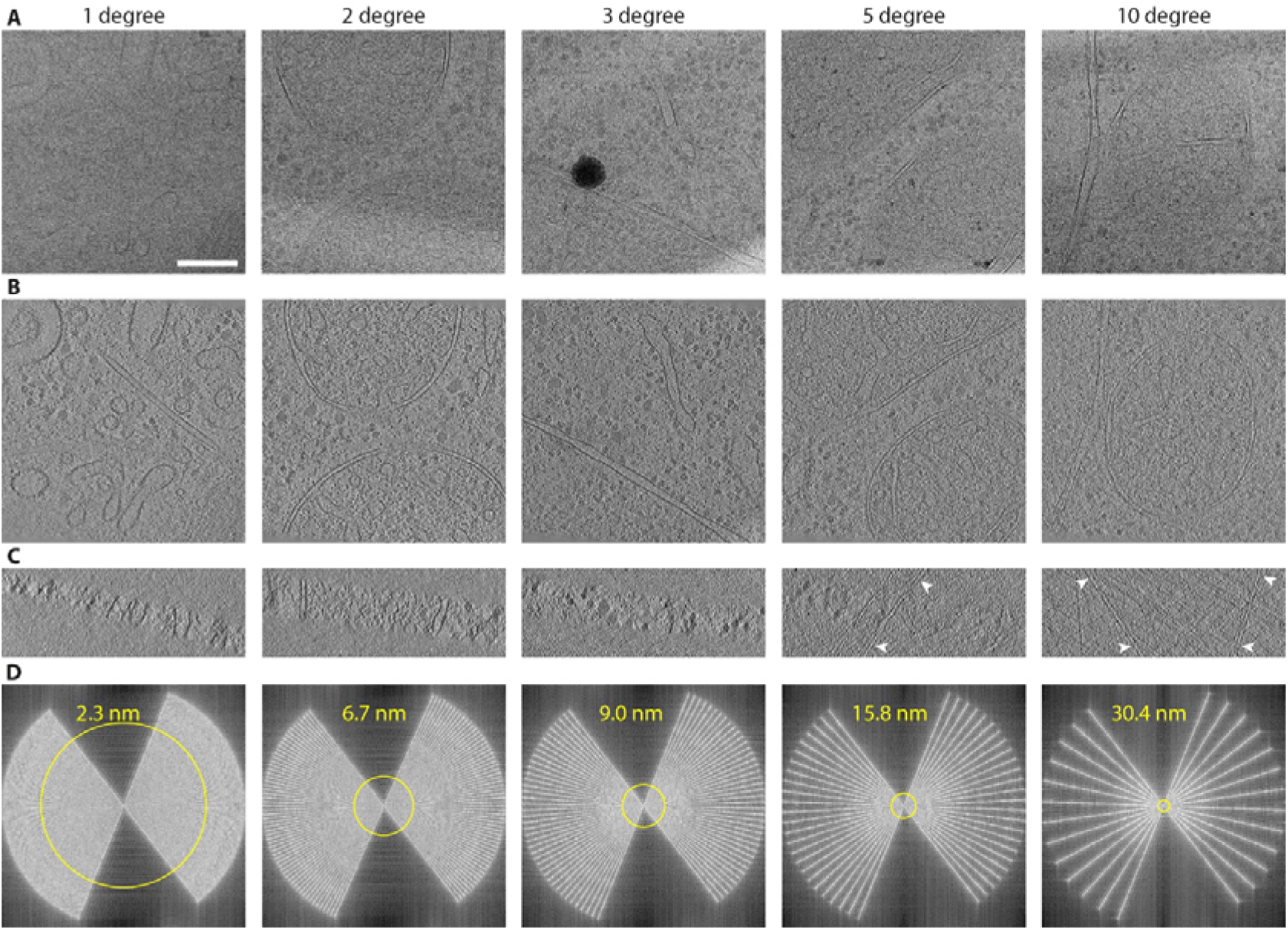
Representative tilt images and tomograms at varying tilt increment. **A** Single projection images of the sample at effective zero-tilt from a tilt series acquired with a tilt increment of: 1 degree (i); 2 degrees (ii); 3 degrees (iii); 5 degrees (iv); and 10 degrees (v). **B** Central XY slices through the tomograms reconstructed from the tilt series shown in A. **C** Central XZ slices through the tomograms shown in B. White arrowheads point to membranes that appear to extend outside of the lamella. **D** Power spectra of representative tomograms for all five conditions. Yellow circles represent the Crowther criterion for the tomograms shown in B and C, with the numerical value shown as text in yellow. Scalebar is 200 nm and applies to all relevant panels.

**Table 1.**
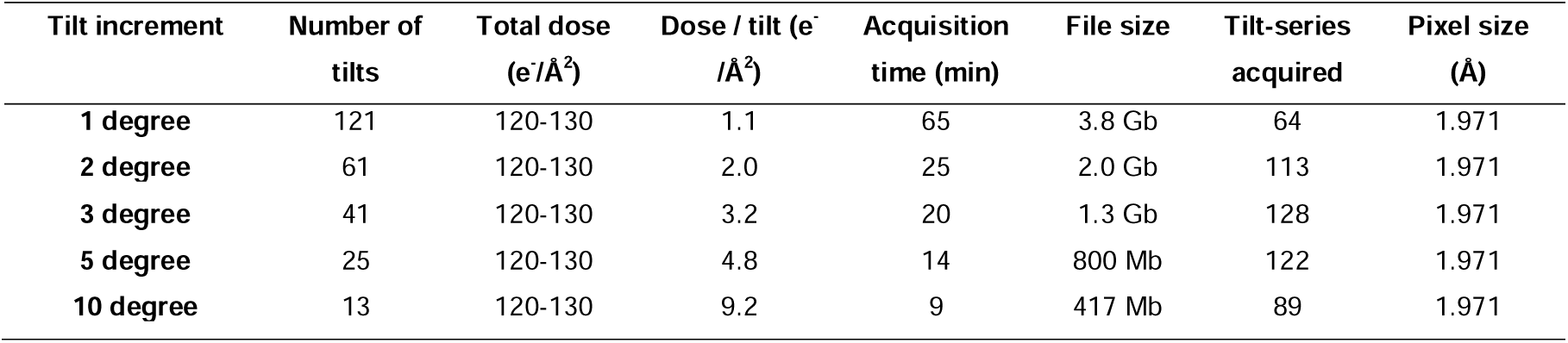
Datasets and acquisition parameters used in this study.

### Reconstructions visually suffer from higher increments

Representative tilt images, slices through reconstructions, and the respective Crowther criteria are visualized in Fig. 1, SFig. 2 and Supplementary Movies 1-10. All tomograms were successfully reconstructed, even with the unusually large angular increment of 10 degrees. However, in reconstructions obtained from the 5- and 10-degree tilt-increment data, membranes were observed to extend outside of the lamella body (white arrowheads in Fig. 1C). Furthermore, a loss in tomogram contrast that was primarily noticeable in the XZ-slices through the volumes was apparent (Fig. 1C). In particular, tomograms with 10-degree increment showed severe artefacts surrounding highly curved membranes, and membranes extending far outside of the lamellae (see SFig. 3). The Crowther criterion (Fig. 1D) extended from 2.3 nm in the 1-degree data, to 30.4 nm in the 10-degree data, but also depended on the local lamella thickness in the individual tomogram.

**Fig. 2.**
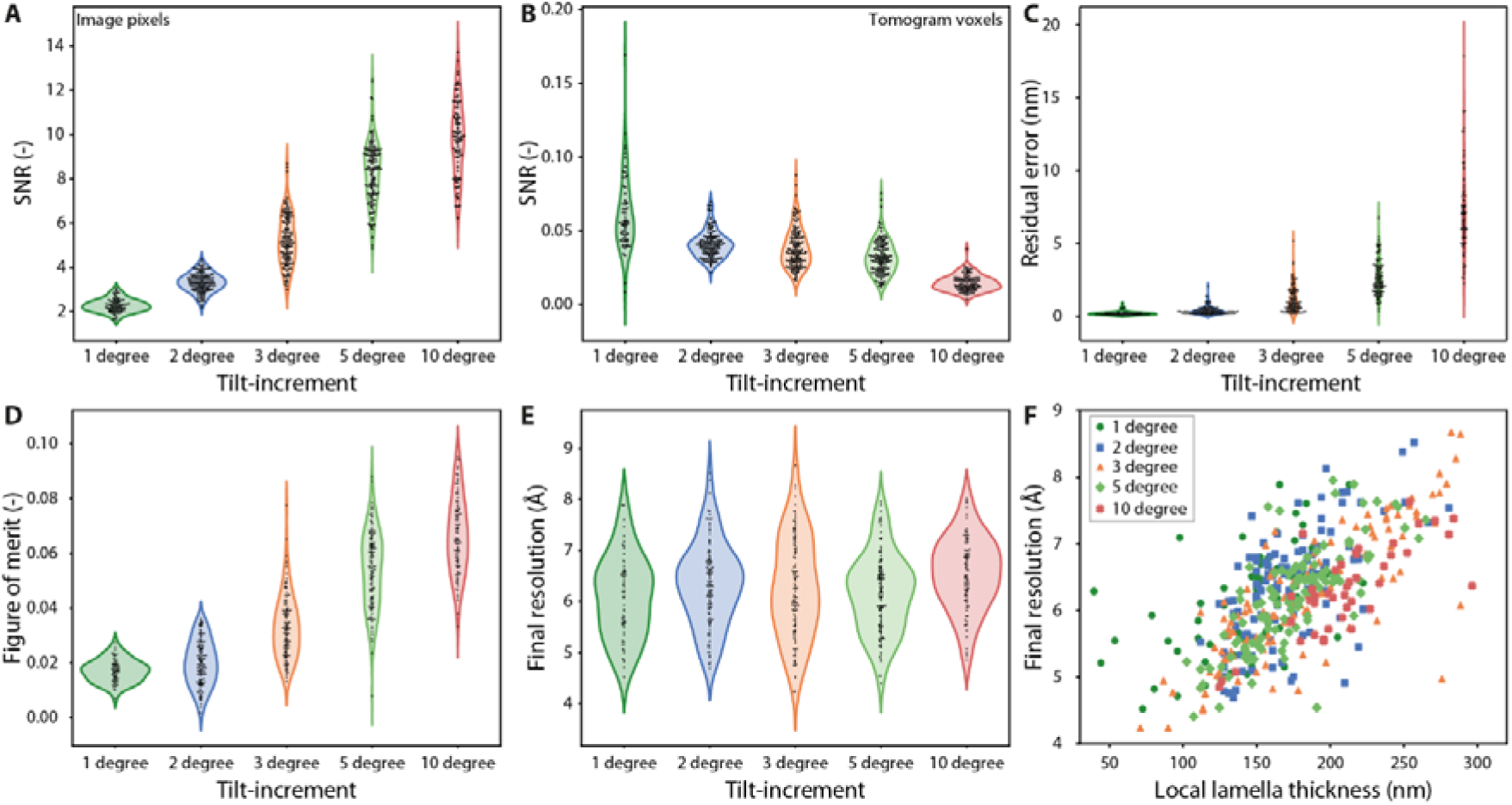
Assessment of signal-to-noise ratio (SNR), tilt-alignment quality and CTF fitting parameters at different tilt increments. **A** SNR measured from projection images of the sample at effective zero-tilt position. SNR was defined as the ratio of the squared mean to the squared standard deviation of each image. **B** SNR measured from the sample volume that represents the lamellae. **C** Residual alignment error after tilt-series alignment in Etomo. **D-E** CTF estimation parameters. **(D)** Figure of merit, which indicates the confidence level to which the CTF was fitted, and **(E**) the final resolution to which the CTF was fitted; both estimated with Gctf on projection images of the sample at effective zero-tilt position. **F** Scatter plot showing the final resolution as shown in (E) against the local lamella thickness for all conditions, with coefficients of determination of R^2^ = 0.30, 0.38, 0.66, 0.61 and 0.60 for the respective datasets.

**Fig. 3.**
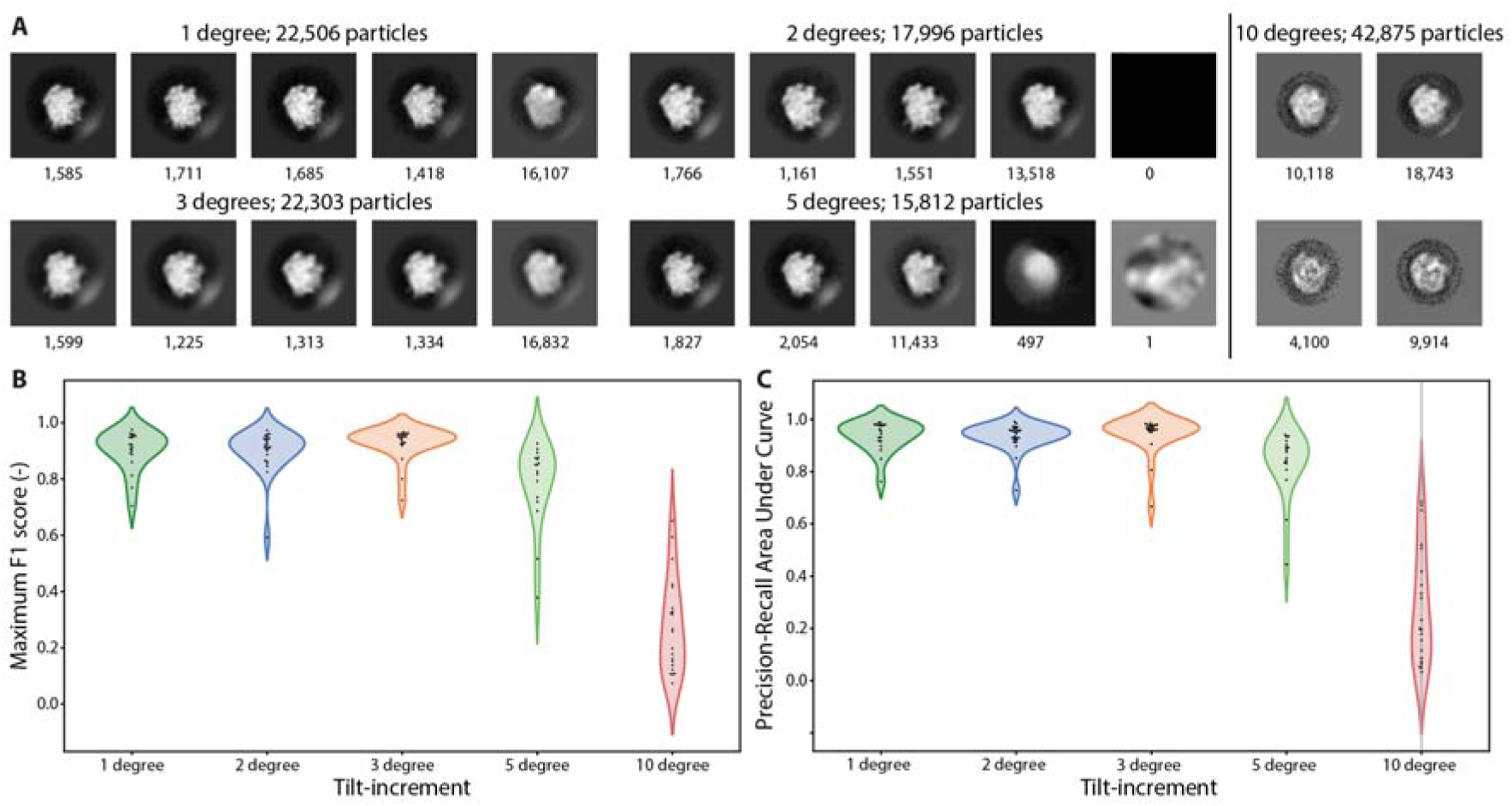
The effect of tilt increment on template matching. **A** Classes after the first round of 3D classification. **B** Maximum F1-score for tomograms used for TM and STA. **C** Area under the precision-recall curves. See corresponding SFig. 9.

### The residual alignment error suffers from higher increments

Next, we quantified the SNR of the tilt images and the tomographic volumes by dividing the square of the mean by the square of the standard deviation of the images (for details, see Methods). As expected, due to the increased dose per image, the SNR in tilt images increases with the increment (see Fig. 2A). However, how the SNR of projection images affects the reconstructions further depends on the alignment accuracy. When analysing tomographic volumes, we found that tomograms from data with a smaller increment displayed higher SNR values (see Fig. 2B), whilst showing a similar lamella thickness distribution (see SFig. 4A). This is also apparent in the XZ-slices shown in Fig. 1C. An insightful metric to judge the accuracy of the alignment of projection images is the residual error that is calculated by the Etomo program in IMOD (*17*), which is the deviation of tilt patches from the alignment fit during tilt-series alignment (*17*). Interestingly, large differences were observed for this metric when comparing the datasets (Fig. 2C). We found that tilt series with a smaller tilt increment could be aligned with higher accuracy. These data show that tilt images with as little as 1.1 e^-^/Å^2^ were already aligned with great accuracy.

**Fig. 4.**
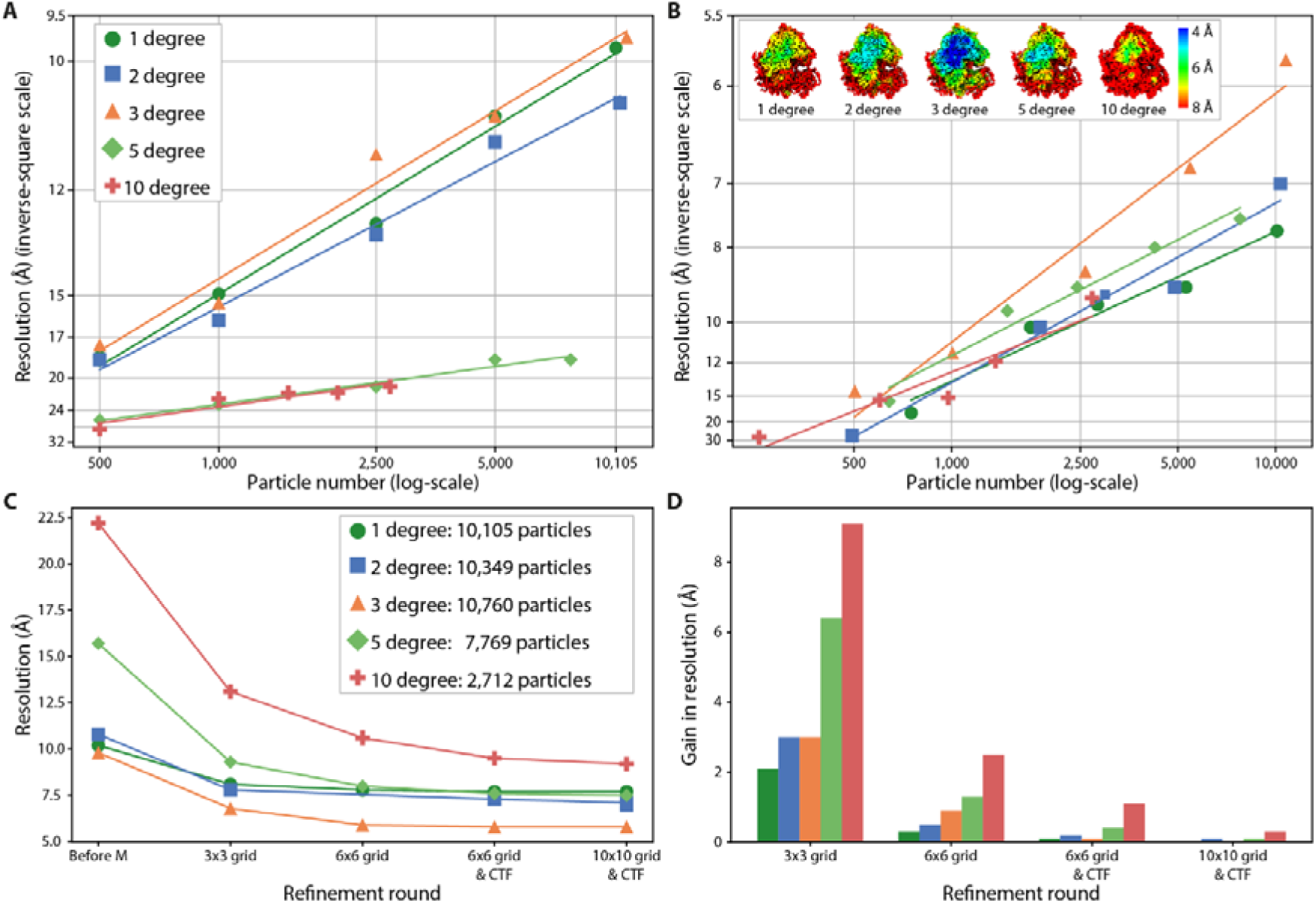
The effect of tilt increment on STA. **A** Rosenthal–Henderson plots of particle subsets when refined and averaged in RELION 3.1. **B** Rosenthal–Henderson plots of tomogram subsets when refined and averaged in M. Inset: ribosome density maps using all particles, coloured according to the local resolution. **C** Resolution progression for each M refinement, considering all available particles. The grid subdivision shown on the horizontal axis is used for both the Image warp and the Volume warp grids. For more details, see the Methods section. **D** Resolution gain for each refinement round, also shown in panel C.

After estimation of the contrast-transfer-function (CTF) using Gctf (*40*), we found that the CTF was fitted with higher confidence to images with higher dose based on the metrics “figure of merit” and “CC-score”, as expected. Surprisingly though, the resolution to which the CTF was fitted was similar for all conditions, despite an 8-fold increase in dose (see Fig. 2E). Similar results were obtained using CTFFIND4 (*41*) (SFig. 5). In cryo-ET, several factors limit the resolution up to which CTF can be reliably estimated. This includes the dose per image and sample thickness determined by the lamellae, as well as tilt-induced effects such as defocus gradients and increased sample thickness. As the data shown in Fig. 2D-F pertains to images of the untilted specimen, the tilt-induced effects can be ruled out. Moreover, the results in Fig. 2E demonstrate that the observed maximum resolution, at least under the chosen experimental conditions, is not constrained by the per-image-dose. We therefore examined whether the maximum resolution depends on the local lamella thickness in each tomogram. Indeed, the variation in maximum resolution correlates with lamella thickness across all datasets (see Fig. 2F).

From these data, we conclude that tilt images can be acquired and aligned for all conditions, but smaller tilt increments result in tomograms that are better aligned and have a higher SNR.

### Template matching performance suffers from higher increments

Ribosomes are widely abundant throughout the cytoplasm and, due to their size and electron contrast, have a high degree of processability. We therefore used ribosomes as probes to study the effect of tilt increment on TM and STA, as was done previously to study other effects (*31*, *42*). Firstly, we selected ca. 20 tomograms per condition, based on tomogram content and local lamella thickness (*31*) (for more details, see Methods and SFig. 4B). We then performed TM for ribosomes on the selected tomograms with a binning factor of 4 (corresponding to a voxel size of 7.9 Å), and an angular search of 5° using GAPSTOP^TM^ (*24*, *31*, *43*) (see SFig. 6). For the 1-, 2- and 3-degree-increment data, we could set peak extraction thresholds such that only true ribosomes were selected, and no 3D classification was necessary to remove, or able to detect junk particles (see SFig. 7). For 5- and 10-degree data, however, 3D classification in RELION 3.1 (*44*) had to be utilised to separate junk particles from the TM results (see SFig. 8). For the 10-degree data, 3D classification did not work reliably with a binning factor of 6 that was sufficient for the 5-degree data. Instead, we used a binning factor of 2, still with limited success (see SFig. 8). Finally, in a similar number of tomograms, we identified fewer ribosomes for the 5- and 10-degree data (see SFig. 8). To ensure similar data processing strategies for all conditions, extraction thresholds were also lowered for the 1-, 2- and 3-degree conditions, and 3D classification was performed to filter out the junk particles (see Fig. 3 A and SFig. 8).

By comparing particle lists from before and after 3D classification, we calculated F1-scores and precision-recall (PR) curves for each tomogram used for TM (see SFig. 9 and Methods). To compare all tomograms from each dataset, we plotted the maximum F1-score, and calculated the area under the PR curves for each tomogram (Fig. 3B and C). In summary, TM performed equally well on tomograms from the 1-, 2-, and 3-degree increment data. In contrast, performance was reduced for the 5-degree and even more so for the 10-degree data (Fig. 3B and C).

### Per particle refinement using STA has an optimal tilt increment

We then proceeded with STA, following the well-established Warp-RELION-M pipeline (*32*, *44*, *45*). Briefly, tilt images and tilt-series alignment files from Etomo were imported into Warp for CTF-estimation, tomogram reconstruction, and subtomogram extraction. Initial particle refinement in RELION 3.1 served as the starting point for subsequent multi-particle refinement in M (see Methods). We found that RELION 3.1, which treats the subtomograms as static 3D volumes, refined 1-, 2- and 3-degree data to roughly the same resolution (10 – 11 Å), while the 5- and 10-degree data only averaged up to 19-23 Å (see Fig. 4A and SFig. 10A). After subsequent M refinement, the 3-degree data showed a markedly higher resolution than all other conditions, averaging to 5.8 Å, compared to 7.0 Å and 7.7 Å for the 1- and 2-degree data respectively (see Fig. 4B and SFig. 10B). Furthermore, M improved the resolution for the 5- and 10-degree data to 7.5 Å and 9.2 Å respectively, which is very similar to the 1- and 2-degree data when adjusted for particle numbers (see Fig. 4B).

For each condition, all particles were processed once more in subgroups of different sizes to produce Rosenthal–Henderson plots (*46*), that capture the dependency of resolution on particle number (Fig. 4A & B). To capitalise on M’s multi-particle refinement framework, subgroups were defined based on the selection of tomograms rather than an exact number of randomly chosen particles, as was done for the RELION results.

To investigate the results from M further, we plotted the reported resolution through the various rounds of refinement, as shown in Fig. 4C, D. We found that the resolution improvements per iteration correlated with the amount of dose in the tilt images.

From our STA-analysis, we conclude that, when subtomograms were treated as independent and static 3D volumes, 1-, 2-, and 3-degree data averaged to similar resolution. However, the per-particle tilt image refinement from M yielded the highest final resolution for the 3-degree tilt increment data.

## Discussion

Cryo-ET has recently matured into a robust and accessible technique, supported by advances such as improved hardware, fast parallel acquisition (*47–49*), and streamlined data processing routines. As a result, it is becoming a mainstream tool for *in situ* structural biology. Strategies to optimise data acquisition have thus come into focus, and more systematic research has been performed on how parameters such as lamella thickness and acquisition schemes influence the resolution attainable with STA (*16*, *31*, *42*). In this study, we systematically investigated the effects of varying the tilt increment when collecting cryo-ET data, with the total dose and tilt range kept constant. We found that finer increments (1 – 3 degrees) showed overall similar results in visual tomogram quality, tilt-series alignment, high-confidence TM and STA resolution, whereas coarser increments (5 and 10 degrees) compromised tomogram quality, showed reduced TM accuracy, and subsequently lower STA-resolution. However, after M refinement, the 3-degree dataset clearly outperformed all other datasets in terms of resolution. For STA, a 3-degree increment may provide the optimal balance between angular sampling, dose distribution, and computational expense during processing.

To further contextualise these findings, we first examined the quality of the tomograms. Contrary to our expectations, angular sampling, rather than the amount of dose per projection image, was the primary factor that influenced the alignment accuracy as approximated by the residual error. This observation may be explained with differences in alignment strategies used for *in vitro* and *in situ* samples. For *in vitro* studies, gold fiducial beads are typically used for alignment. These beads retain their spherical shape and similarity, regardless of the projection angle or the angular gap between successive images, thereby providing consistent and robust reference points across the tilt series. In contrast, alignment of *in situ* datasets often relies on patch tracking of endogenous structural features, which are generally irregular in shape. Their projection geometry varies with the projection angle and they therefore lose similarity between successive images more rapidly. As a result, the cross-correlation peaks become broader or less well-defined, ultimately reducing alignment accuracy.

We further found that using a 1 – 3-degree increment, despite the reduced SNR in the tilt images, generally led to high contrast in the tomograms. Initially, we reasoned that finer angular sampling would extend the Crowther criterion to higher spatial frequencies and, when combined with improved alignment, would yield tomograms of visibly enhanced quality. However, we could not visually distinguish tomograms coming from 1-, 2-, or a 3-degree tilt series. Conversely, coarser increments (5- and 10-degree) compromised tomogram quality and SNR and led to artefacts in the tomograms. Furthermore, whilst the 1-degree data exhibited the best alignment, it is associated with practical drawbacks. For instance, the acquisition time per tilt series increased more than 2-fold compared to 2-degree tilt increments. In addition, a higher number of projections increases demands on data storage and computational resources during pre-processing, including subsequent steps that utilise tilt images or intermediate data. However, differences in storage and processing requirements vanish once actual 3D tomograms or subtomograms are reconstructed or extracted, but re-emerge during subsequent M refinements.

As cryo-ET datasets are becoming increasingly larger, and also publicly available, the focus in the field shifts from data acquisition to accurately and efficiently mining the wealth of information that is present in existing cryo-ET datasets (*50–54*). TM has shown itself to be a robust and sensitive tool to detect the structural signature of proteins (*24*, *55*), but is computationally very expensive, which has become a major bottleneck of the workflow in practice (*56*). Moreover, further particle curation is often necessary to remove false-positive TM picks from the true-positive particles through extensive 3D classification procedures. It is therefore important to understand how the tilt increment influences the TM procedure to enable its optimisation. Here, we found that TM performed equally well for the 1-, 2-, and 3-degree increment data on ribosomes, and peaks higher than a threshold could be extracted to yield only true positives so that successive 3D classification was not necessary. This was not possible for the 5- and 10-degree data, however, likely due to a combination of suboptimal tilt-series alignment and reduced SNR in the tomograms. We hypothesise that for smaller templates that comprise more challenging targets, improved tomogram alignment and increased Crowther criteria are crucial for successful TM, thereby favouring the usage of smaller tilt increments, particularly when TM is performed on data with a lower binning factor. This notion is supported by recent work, that showed that the addition of interpolated tilts, generated using cryoTIGER, improved the performance of TM of nucleosomes (*57*). We anticipate that a well-balanced choice of experimental angular increments and tilt interpolation during post processing may be key to future workflows that will benefit from optimal angular sampling.

In STA, the incomplete angular sampling of individual particles may be less critical compared to the analysis of entire tomograms and TM, due to the large number of particles contributing to the final reconstruction. The initially limited angular sampling can be compensated by the inclusion of particles with slightly different orientations, provided that a sufficient number of particles is available. Furthermore, for datasets with fewer projections, suboptimal initial alignment may be mitigated by the higher SNR of individual tilt images. When analysing STA results in our data, as with TM, we could not find major differences in the 1-, 2-, and 3-degree datasets when averaged with RELION 3.1, whereas the 5- and 10-degree datasets yielded a noticeably lower resolution. Using M improved the resolution across all datasets to comparable levels, accounting for reduced particle numbers, and notably rescued the 5- and 10-degree datasets despite their lower initial resolution. We hypothesise that two factors contribute to the pronounced tilt increment dependency of the resolution gain after M. Firstly, the initial tilt-series alignment was markedly poorer at higher increments, leaving more potential for improvement. Secondly, these images were acquired at a higher dose, and therefore have a higher SNR, which likely facilitates more effective alignment by M.

Remarkably, the 3-degree dataset reached a higher resolution than all others. Acquiring data with a 3-degree tilt increment likely strikes an optimal balance for the spatial refinement of projections on a per-particle basis STA refinement: it enables robust initial tilt-series alignment and high SNR in the tomograms to support successful TM, while still delivering sufficient dose per tilt image to facilitate effective M refinement. Furthermore, especially compared to the 1-degree data, data collection with a 3-degree increment is more streamlined in terms of acquisition time and data storage.

We note that, while we believe our findings can be applied to cryo-ET in general, the present investigation was confined to a single sample and molecular target. Additionally, although the exact STA resolution may vary, the general trends identified here are unlikely to change when different software packages are used.

Furthermore, we focused our research on *in situ* data. It is unclear to which extent these results are transferrable to *in vitro* cryo-ET studies of inherently thin samples. The alignment of those tilt series is usually performed using gold fiducial beads, which provide great precision, as discussed above. Furthermore, reduced tilt ranges may be used to increase signal in the lower tilt images, however, alignment will most likely not be affected.

Ribosomes are abundant, electron dense, and the processing workflow is particularly effective for ribosomes, making them ideal candidates for studies of this nature. However, it remains unclear whether other targets such as smaller particles or particles with a lower abundance, would follow a similar trend, in particular with respect to TM and M refinements. M allows the simultaneous refinement of multiple molecular species within a single dataset. As a result, lower-abundance targets may benefit from alignment driven by nearby ribosomes, provided there is sufficient spatial proximity, and from the advantages of the multi-particle refinement framework. Many *in situ* cryo-ET studies focus on targets at least partially situated within the cytoplasm, which is filled with relatively dense structures and membranes, thereby aiding tilt-series alignment. Large vacuoles, the extracellular space, or the nucleoplasm, where these contrast-rich structures are absent, may comprise more challenging environments. To what extent tomogram content affects tilt-series alignment remains to be further investigated.

Taken together, our findings support the use of a 3-degree tilt increment for tilt-series acquisition, as it yields tomograms with high SNR, accurate tilt-series alignment, robust performance during TM, and maximised resolution when using multi-particle refinement in M. From our perspective, a 3-degree increment also represents an economical choice, reducing beam time through shorter acquisition, and lowering data storage and computational demands. Smaller or more challenging targets may still benefit from finer increments.

## Methods

### Cryo-ET sample preparation

*D.discoideum cells* (strain Ax2-214) were grown in HL5 medium (Formedium) containing 50 µg/mL ampicillin and 20 µg/mL geneticin G418 (Sigma Aldrich) at ca. 20 °C until exponential growth was achieved.

EM grids, R1/4 200 mesh Au grids with SiO2 support film (Quantifoil), were glow discharged for 90 sec at 0.38 mbar and 15 mA using an easyGlow instrument (PELCO) immediately before usage. The cells were diluted to a concentration of 3.3 x10^5^ cells/ml, and a droplet of 100 µl cell suspension was placed on each grid, after which the cells were allowed to attach to the grid for 2-4 hrs at room temperature in the dark. The grids were subsequently blotted and vitrified by plunge freezing into liquid ethane using a Leica GP2 plunger. Cryo-FIB milling was performed using an Aquilos cryo-FIB system (Thermo Scientific). Prior to gallium FIB-milling, grids were coated with an organometallic platinum layer using a gas injection system for 10 sec and additionally sputter coated with platinum at 1kV and 10 mA current for 20 sec. SEM imaging was performed at 10 kV and 13 pA current to guide the milling progress. Milling was performed at 30 kV in a stepwise fashion, where the current was reduced from 500 pA to 30 pA whilst reducing the thickness of the remaining lamella. Finally, lamella polishing was performed with 30 pA current.

### Cryo-ET acquisition

Cryo-ET datasets were collected at 300 kV on a Titan Krios G4 microscope equipped with a cold FEG, Selectris X imaging filter, and Falcon 4 direct electron detector, operated in counting mode (all from Thermo Scientific). Medium magnification overview montages of lamellae were acquired with 3.0 nm pixel size to inspect the lamellae and determine positions for tilt series collection. Tilt series were acquired using SerialEM (versions 4.0.6, 4.0.10, and 4.0.20) in low-dose mode as dose-fractionated movies with size 4096 x 4096 at a magnification of 64,000x, corresponding to a calibrated pixel size of 1.971 Å. Dose-fractionation and motion-correction were performed on-the-fly using 10 equally dosed fractions per tilt movie in SerialEM. Tilt series acquisition started from the lamella pretilt of +8°, and a dose symmetric acquisition scheme was applied (*11*), with tilt increment and exposure as described in Table 1, all with a tilt grouping of 2. The energy filter was set to a slit width of 10 eV, and a nominal defocus of −2.5 to −5 µm was used. The dose rate on the detector was targeted to be ca. 6 e^-^/px/s at a representative spot on the specimen.

### Tomogram reconstruction

The tilt series were dose-filtered using custom Matlab scripts (*58*) and manually cleaned based on visual inspection of the tilt images to remove unusable projection images. CTF-estimation was performed on the tilt images using Gctf (*40*) and CTFFIND4 (*41*). The dose-filtered tilt series were then aligned through patch-tracking in IMOD 4.11.5 (*59*) and reconstructed using weighted back projection and fSIRT for visual inspection. During fiducial model generation using tiltxcorr, optimal filter parameters were found for several representative tilt-series for each dataset, after which these were applied to all tilt-series in the dataset. Specifically, low-frequency rollof sigma was 0.01 for the 1-and 3-degree datasets, 0.02 for the 2-degree dataset and 0.001 for 5- and 10-degree datasets. High frequency cutoff radius was 0.1 for the 2-degree data, and 0.08 for all other datasets. High frequency rolloff sigma was kept constant at 0.05. During tomogram reconstruction, all images were multiplied with a factor of 250 by the tilt command. To ensure compatibility with the Warp and M software suites, tomograms were then reconstructed again in Warp, using the alignment obtained from Etomo. Both standard and deconvolved tomograms were reconstructed, using Warp’s standard tomogram reconstruction settings that weighs and corrects the data for CTF, dose and tilt. Local lamella thickness was measured manually for each tomogram using IMOD.

### Crowther criterion

The Crowther criterion, or the resolution to which the tomogram is completely filled (disregarding the missing wedge), *d* is given by:

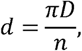

where *n* is the number of projection images and *D* is the specimen thickness (*19*). The conversion of resolution to number of pixels in the Fourier transform was done using the function resolution2pixels from cryoCAT (*60*). When calculating the numerical values for the Crowther criterion, we simplified by assuming evenly spaced projections over the full tilt range, thereby disregarding the missing wedge.

### SNR measurements

The SNR was calculated according to:

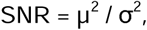

with *µ* the mean of the image and σ the standard deviation of the image (*61*). For tilt images, SNR calculations were based on unbinned, unprocessed motion-corrected tilt images of the specimen at effective zero-tilt. In the rare event where this image was removed during pre-processing, the next available tilt image was used instead. For tomogram volumes, voxels outside of the lamella body were masked out, before the SNR was calculated.

To quantify differences in signal-to-noise ratio (SNR) across tilt increment conditions, a non-parametric Kruskal-Wallis test was performed as an omnibus test of the null hypothesis that all groups are drawn from the same distribution. Because SNR distributions were not assumed to be normal, and sample sizes differed across conditions, non-parametric tests were used throughout. Following the omnibus test, all 10 pairwise comparisons between conditions were assessed using two-sided Mann-Whitney U tests. To control for multiple comparisons, raw p-values were adjusted using the Bonferroni correction (multiplied by the number of comparisons, n=10, capped at 1.0). Statistical significance was defined as a Bonferroni-corrected p-value below 0.05. All analyses were performed in Python using the scipy.stats module.”

### Template matching and 3D classification

To ensure processing feasibility, ca. 20 tomograms were selected for each condition to be subjected to TM and STA. Tomograms were selected based on the following criteria: local lamella thickness was limited to ca. 180 nm to optimise resolution (*31*); tomograms that predominantly contained organelles that exclude ribosomes, such as mitochondria or the nucleus, were excluded from analysis. From the resulting list of tomograms, 20 were chosen at random. Care was taken that these tomograms formed a representative subset of the original thickness and alignment distributions to avoid bias by tomogram selection.

TM was performed using GAPSTOP^TM^ (*24*, *62*), a GPU-accelerated implementation of STOPGAP (*43*). As a template, a previously determined high-resolution ribosome map from *D. discoideum* was used, from reference (*31*), and low-pass filtered to a resolution of 30 Å. TM was performed on deconvolved tomograms from Warp with a voxel size of 7.9 Å (corresponding to a binning factor of 4) using an angular search of 5 degrees. TM was performed on a high-performance computing cluster using 16 A100 GPUs (Nvidia), running for approximately 1 hour and 20 min per tomogram, independent from the tilt-increment. The resulting score maps were converted to z-scores for more generalisable peak extraction across tomograms (*24*). Initially, peaks were thresholded using a carefully selected z-score (6, 6, and 5.5 for the 1-, 2-, and 3-degree datasets respectively) to optimise particle extraction (see SFig. 7 and main text). Later, thresholds were lowered to a z-score of 5 for all conditions, except for the 10-degree dataset, where peaks were extracted using a threshold at z-score 3.75. Peak location and angular information from TM were converted to particle STAR files using the cryoCAT package (*60*).

Subtomograms were extracted using Warp at a binning factor of 6 (corresponding to a voxel size of 11.8 Å) and subjected to multiple rounds of 3D classification in RELION 3.1 (*44*) (see SFig. 8) until a clean particle list was obtained.

### Template matching evaluation

To ascertain the quality of the TM procedure, we calculated the precision, recall, and F1-scores. Firstly, ribosome positions confirmed by 3D classification were considered the ground truth (GT). Then, TM peaks were extracted at various z-score thresholds, and these particle locations were compared to the GT. For each extraction, the overlap of GT-particles and extracted particles are true positives (TP). The false positives (FP) are locations that were extracted, but not in the GT. Then, the precision (P) is calculated as:

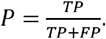

Similarly, the recall (R) is given by:

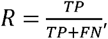

where FN are the false negatives, meaning the particles in the GT list that were not extracted when using that extraction threshold. Precision-recall curves were plotted by calculating both the precision and recall values for all extraction thresholds described above.

The F1 score is the harmonic mean of precision and recall:

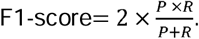

### Ribosome particle refinement

After 3D classification, a 3D refinement was carried out on the particles selected as true positive particles using RELIONs 3D Refine job type. These refined particles were re-extracted in Warp using a binning factor of 2 (corresponding to a voxel size of 3.9 Å), refined again, after which subtomograms were extracted in an unbinned fashion, refined in RELION once more, and the results served as inputs for M (version 1.0.9) (*32*), starting from the on-the-fly frame-aligned projection images.

M refinements were carried out as shown in Table 2. When convergence was reached, no further refinements were carried out.

**Table 2.** Parameters of M refinement rounds.

| M refinement | Geometry |  | Tilt series |  | CTF |  |
| --- | --- | --- | --- | --- | --- | --- |
|  | Image warp grid | Particle poses | Stage angles | Volume warp grid | Defocus | Tilt series acquired |
| Round 1 | 3x3 | ✓ | ✓ | 3x3x2x10 | - | - |
| Round 2 | 6x6 | ✓ | ✓ | 6x6x8x10 | - | - |
| Round 3 | 6x6 | ✓ | ✓ | 6x6x8x10 | ✓ | ✓ |
| Round 4 | 10x10 | ✓ | ✓ | 10x10x10x10 | ✓ | ✓ |

### Rosenthal–Henderson-plots

To produce Rosenthal–Henderson plots (*46*), particle subgroups were randomly selected in two ways. For RELION refinement, the “extract_random” function from starparser was used (*63*) on the unrefined and unbinned STAR file exported from Warp to produce particle subsets of 5000, 2500, 1000, and 500 particles. These particle subgroups were then refined independently using RELION as before, and post-processed using a solvent mask.

For M refinements, particles were pseudo-randomly selected based on whole tomograms using a custom Python script. Care was taken to first process the smallest particle subset in M, and then continue with the subsets containing more particles to avoid cross-interference of the previous refinement rounds.

## Acknowledgements

This work was funded by the Max Planck Society, has been made possible in part by grant number 2021-234666 from the Chan Zuckerberg Initiative DAF, an advised fund of Silicon Valley Community Foundation, and by the Deutsche Forschungsgemeinschaft (DFG, German Research Foundation) – SFB1507 Project No. 450648163 (P17). We thank Patrick C. Hoffmann for help with the cell culture. We thank the Central Electron Microscopy Facility of the Max Planck Institute of Biophysics for providing microscope access and support, and are especially grateful to Mark Linder and Sonja Welsch for technical support. We thank Jan Philipp Kreysing, Sergio Cruz-León, and Gerhard Hummer for useful discussions; Stefanie Böhm and Sonja Welsch for critical reading of the manuscript. We thank Iskander Khusainov, Özkan Yildiz, Juan F. Castillo Hernandez, Andre Schwarz, Erin Schuman, and the Max Planck Computing and Data Facility for support with scientific computing.

## Author contributions

Conceptualization: MWT, BT, MB

Data curation; MWT, MB

Formal analysis: MWT, MB

Funding Acquisition: BT, MB

Investigation: MWT, MB

Methodology: MWT, BT, MB

Project Administration: MWT, MB

Resources: MB

Software: MWT, TM, MB

Supervision: BT, MB

Validation: MWT, MB

Visualization: MWT, MB

Writing—original draft: MWT, MB

Writing—review & editing: MWT, TM, BT, MB

## Data and materials availability

Relevant cryo-ET density maps generated in this study have been deposited in the EM Data Bank (EMDB) with the following access code: EMD-57021. All raw data have been deposited to the electron Microscopy Public image Archive (EMPIAR) database (accessing code EMPIAR-13391), including acquisition metadata, information, tilt-series alignment data, all raw TM locations, and particle locations after 3D classification and refined particle motive lists. All other data needed to evaluate the conclusions in the study are present in the paper.

