## Supplement for "Optimising the tilt-increment for in situ cryo-electron tomography"

Maarten W. Tuijtel^1^, Tomáš Majtner^1^, Beata Turoňová^1^ and Martin Beck^1,2^*,

1 Department of Molecular Sociology, Max Planck Institute of Biophysics, Frankfurt am Main, Germany.

2 Institute of Biochemistry, Goethe University Frankfurt, Frankfurt am Main, Germany

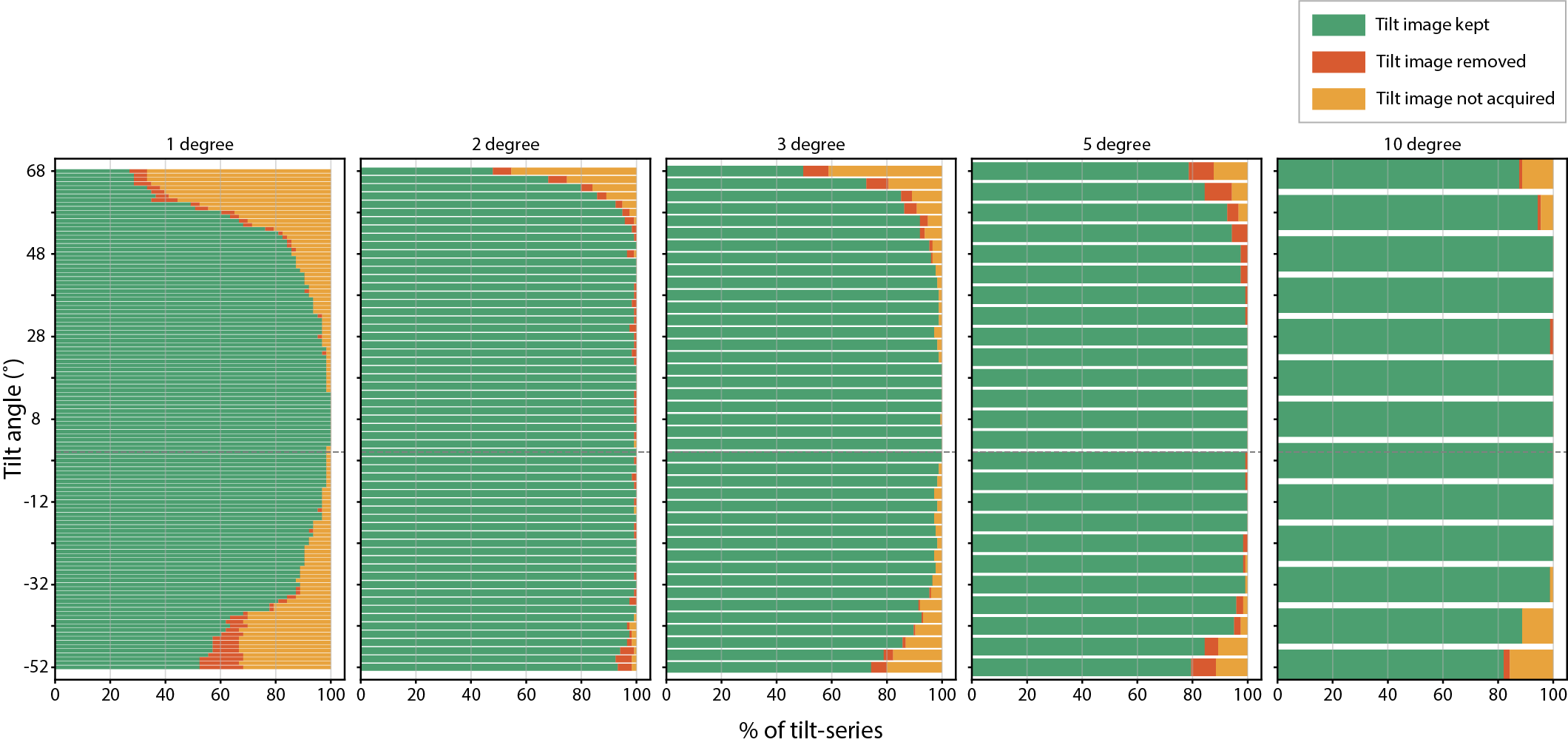

**SFig. 1 Overview of acquired, removed and remaining tilt images for each dataset.** For each dataset, the percentage of tilt images is shown per category: images contributing to the final tomogram (green), images that were removed due to drift, obstruction or other causes (red), and images that were not acquired due SerialEM’s automated abort criteria (orange).

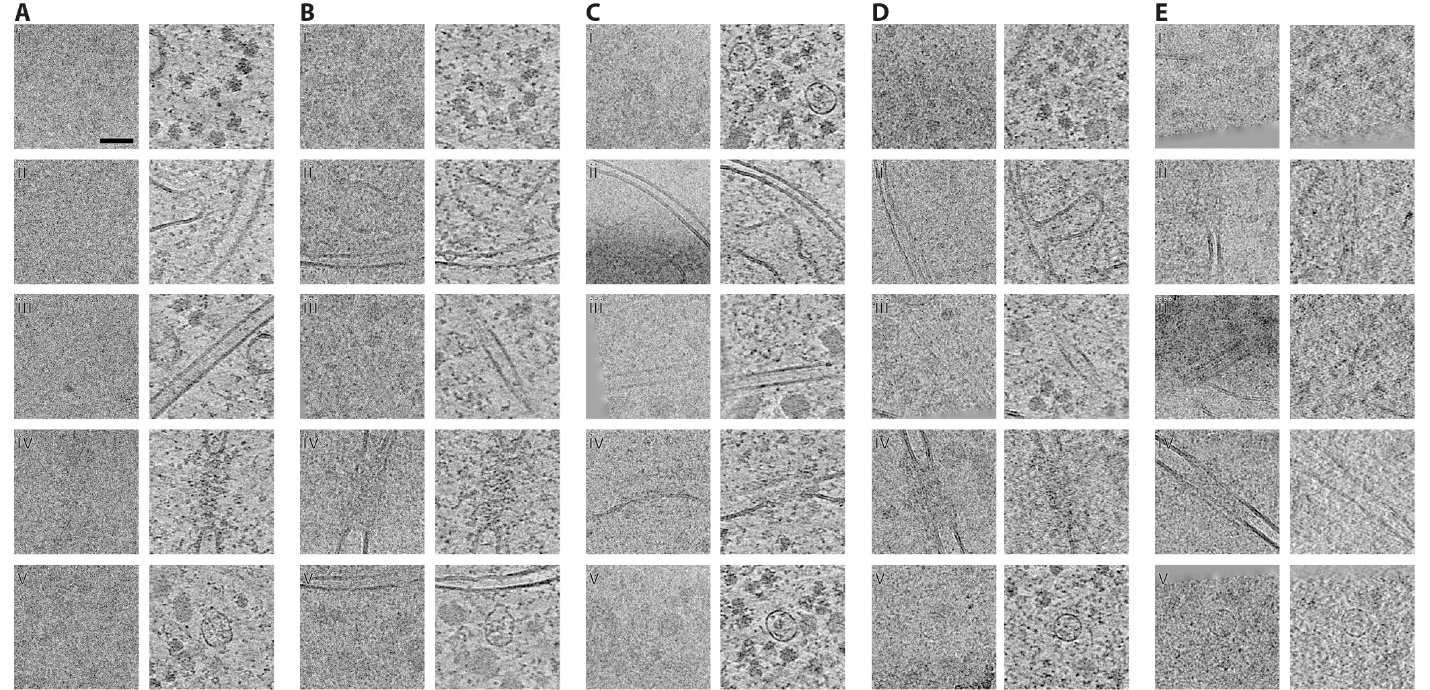

**SFig. 2** **Overview of distinct features in tilt-images and tomogram slices.** **A-E** Columns represent the 1- (**A**), 2- (**B**), 3- (**C**), 5- (**D**) and 10-degree (**E**) datasets. Within each column the central tilt-image is shown on the left and a slice through the tomogram at the same position is shown on the right. The rows represent different features, from top to bottom: (i) ribosomes, (ii) mitochondria outer membrane, (iii) microtubule, (iv) nuclear pore complex, (v) VAULT protein. Scalebar is 50 nm and applies all both panels.

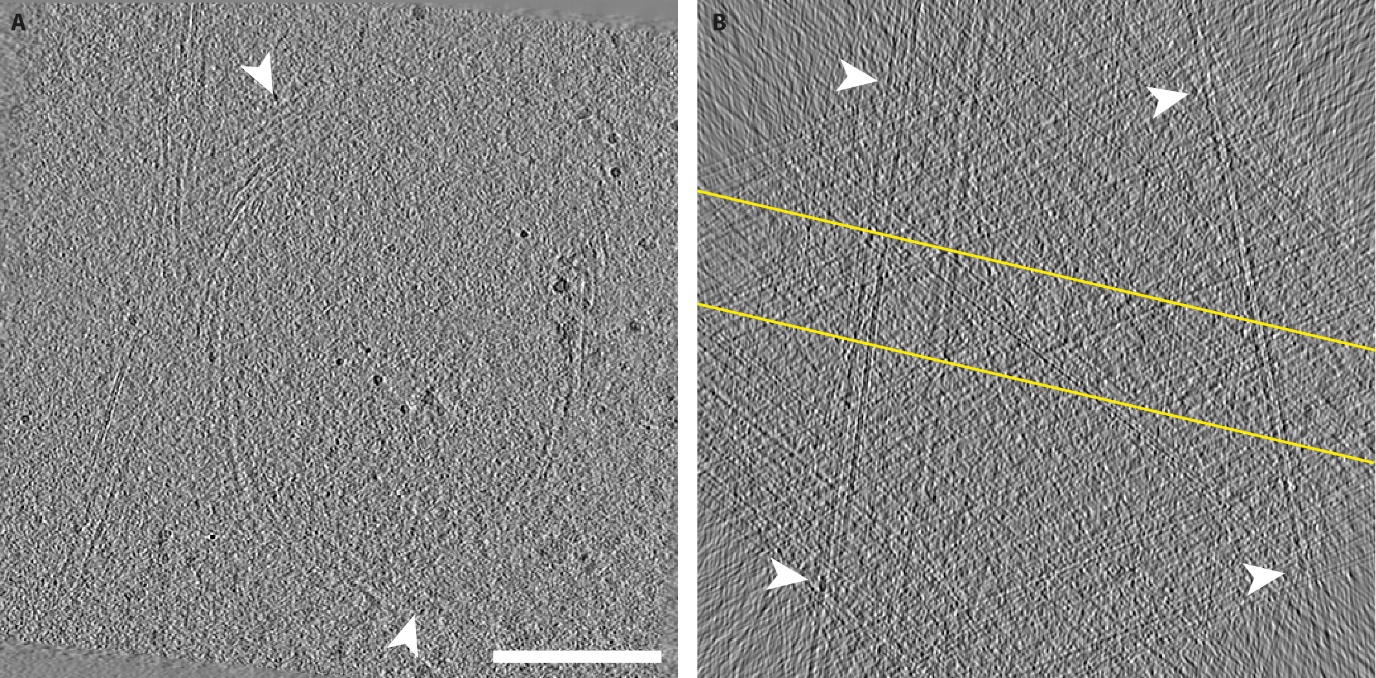

**SFig. 3 Artefacts in tomograms acquired with 10-degree tilt increment.** **A** XY slice of a tomogram, near the lamella surface, reconstructed from a tilt series with a 10-degree increment. Arrowheads point to artefacts in the reconstruction in the region close to strongly curved mitochondrial membranes. **B** Central XZ slice of the same tomogram shown in A, reconstructed with extended Z-height. The region where the lamella is positioned is indicated with yellow lines. Membranes that extend far beyond the lamella are indicated with white arrowheads. Scalebar is 200 nm and applies to both panels.

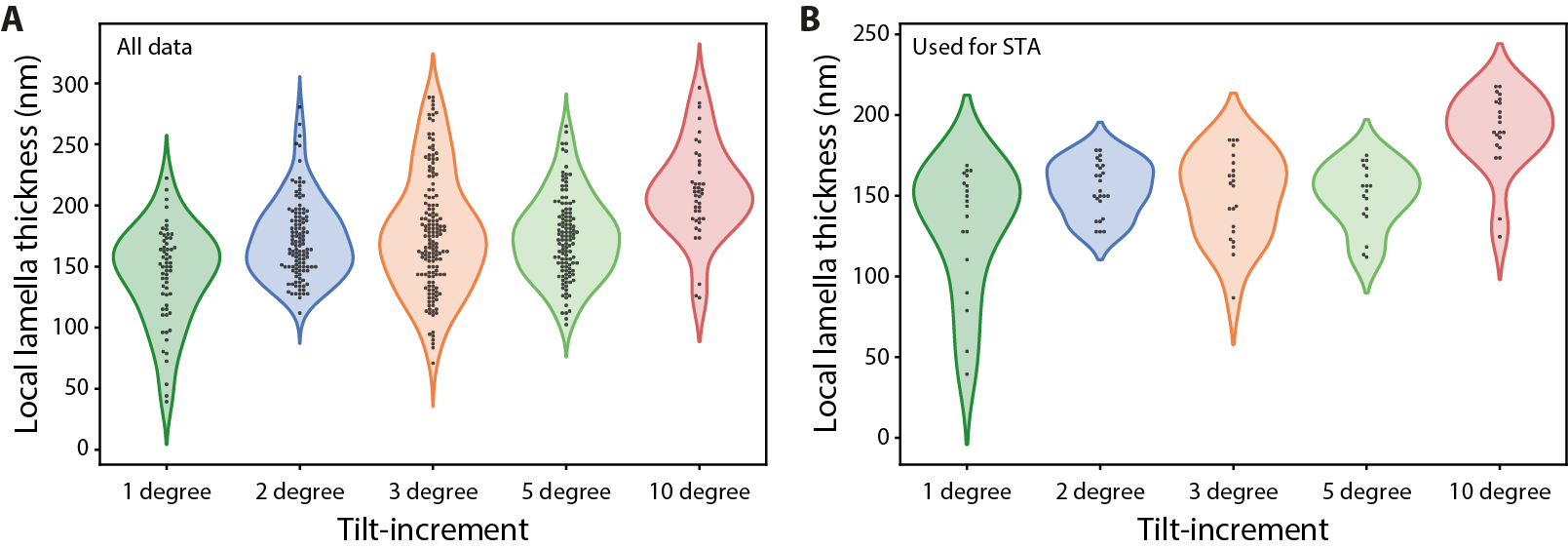

**SFig. 4 Local lamella thickness per condition. A** Local lamella thickness for all tomograms used in this study. **B** Local lamella thickness for all tomograms used for the TM and STA analyses. Owing to the poor tomogram reconstruction quality observed for the 10-degree tilt increment data, lamella thickness could not be determined for each tomogram for this condition.

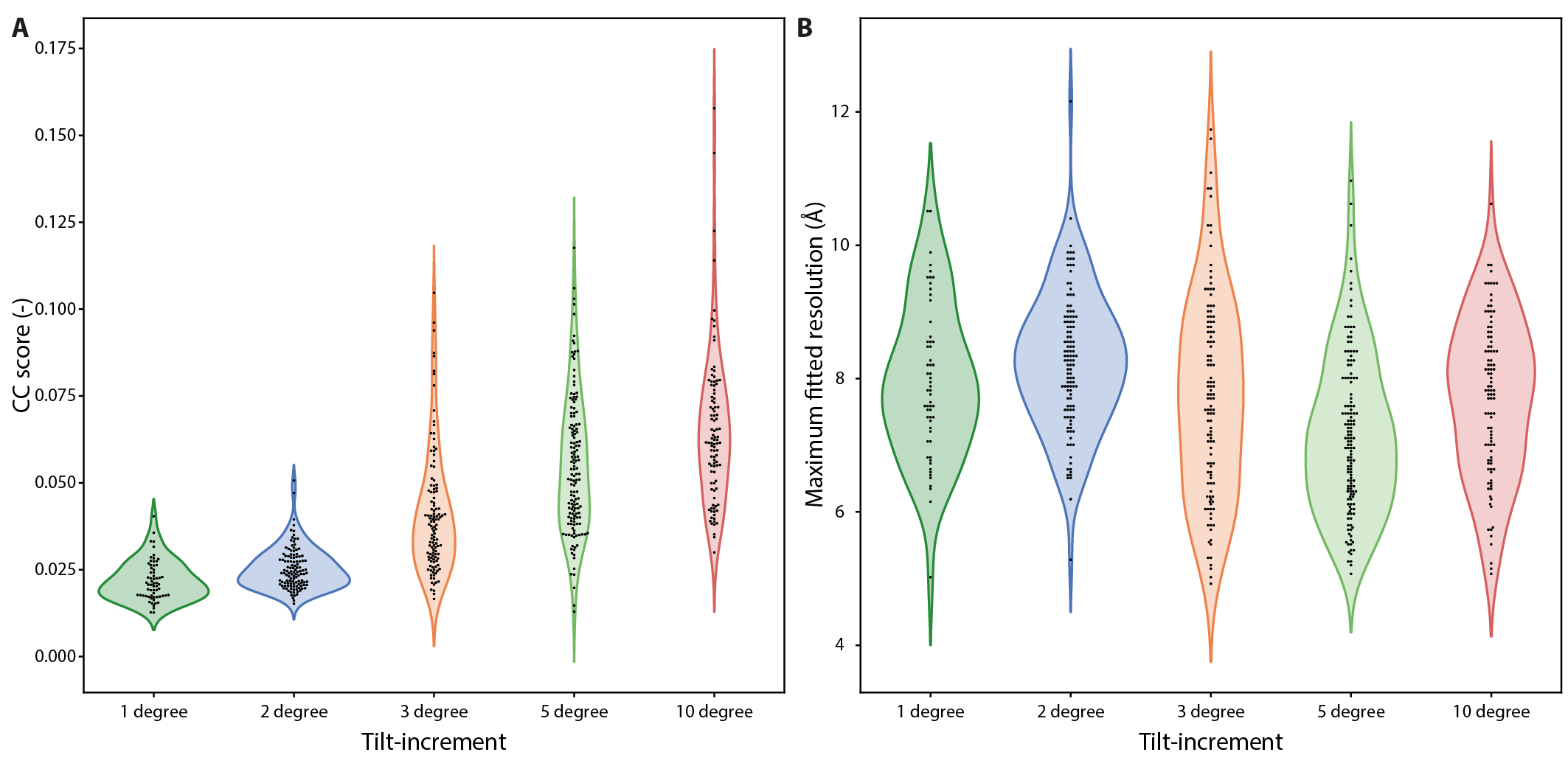

**SFig. 5. CTF estimation parameters from CTFFIND4.** **A** CTFFIND4 CC score, which indicates the confidence level to which the CTF was fitted. **B** Maximum fitted resolution of the CTF using CTFFIND4. For clarity, only the images of the untilted specimen were used in the analysis.

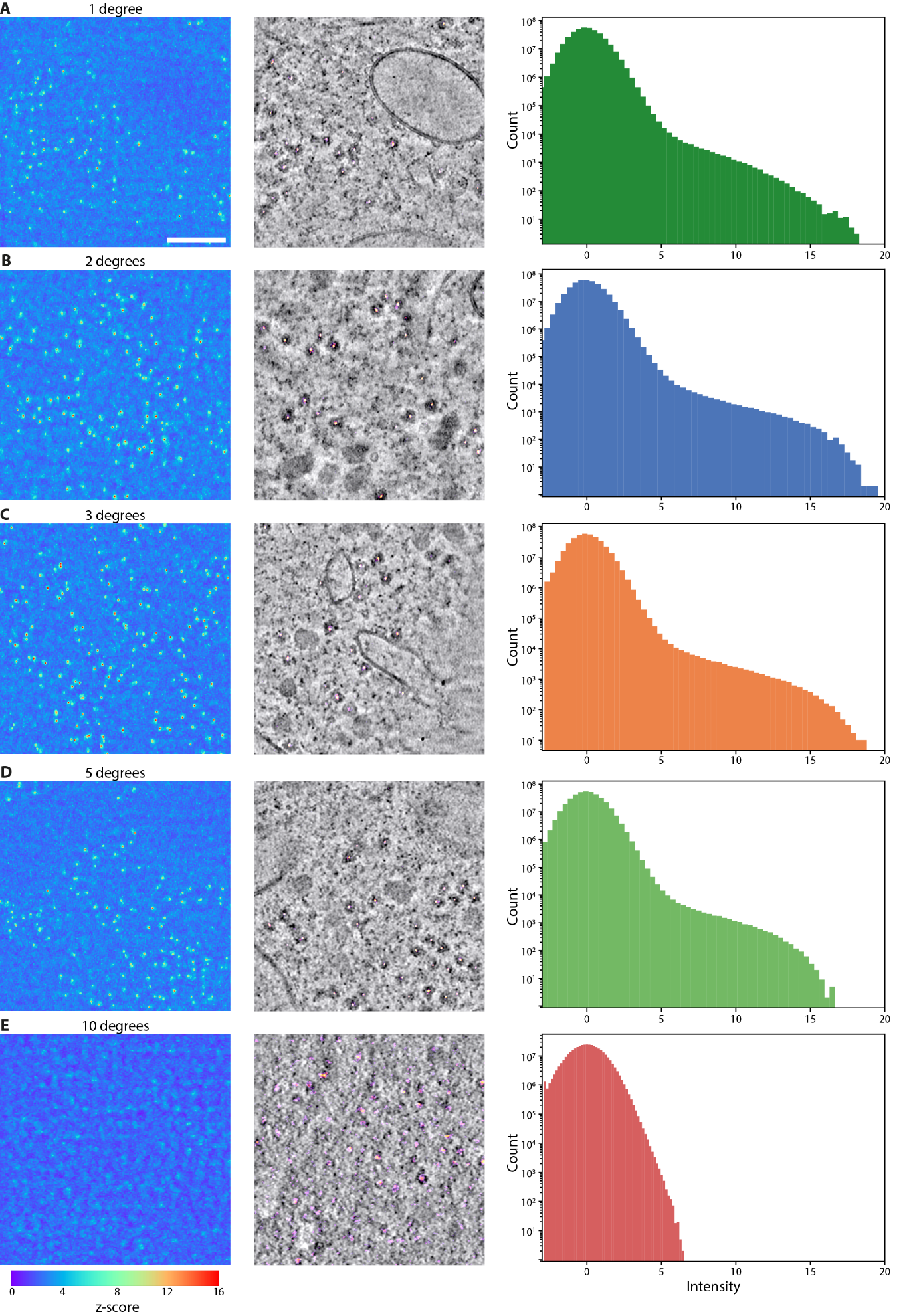

**SFig. 6 Template matching scores**. **A-E** Volumes with template matching scores were converted to z-scores. For visualisation purposes, here only a subvolume of 500 x 500 x 400 voxels were isolated from representative volumes. Panels **A-E** represent results for 1-, 2-, 3-, 5-, and 10-degree increments respectively. A maximum intensity projection was performed along the last dimension (z) and plotted (left). An overlay the scores and the tomogram of the same region (middle) and a histogram of the score volume (right). Scalebar shown in panel A is 100 nm and applies to all relevant panels.

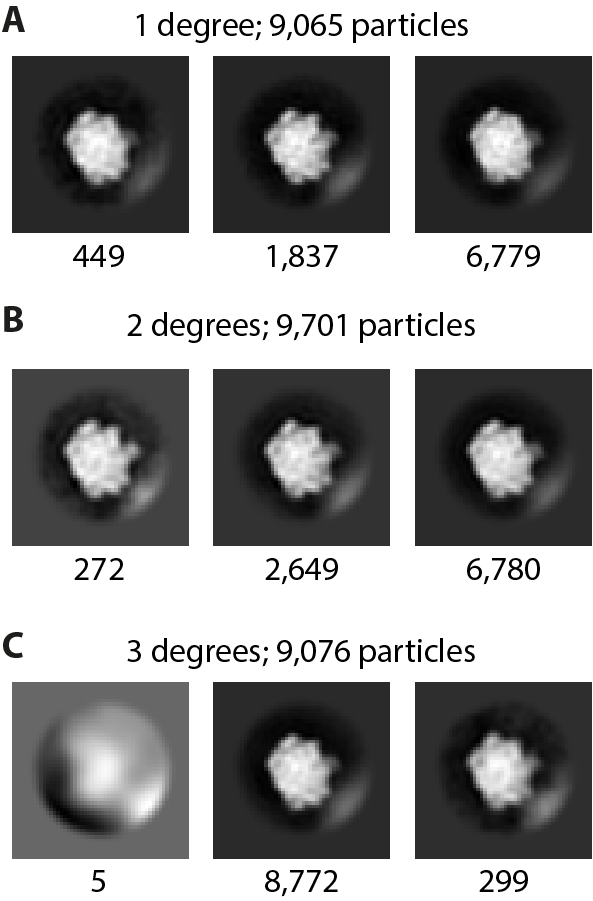

**SFig. 7 3D classification results after high confidence TM.** Resulting classes after 3D classification with the intention to remove junk particles, performed on subtomograms with binning factor 6, corresponding to a voxel size of 11.8 Å, for tilt increments of 1 degree (**A**), 2 degrees (**B**), and 3 degrees (**C**). Z-score thresholds of 6, 6 and 5.5 were used for the 1-, 2- and 3-degree data respectively. The total number of particles is indicated above the classes, as well as the number of particles per class below the class. No significant number of junk particles could be detected for these datasets.

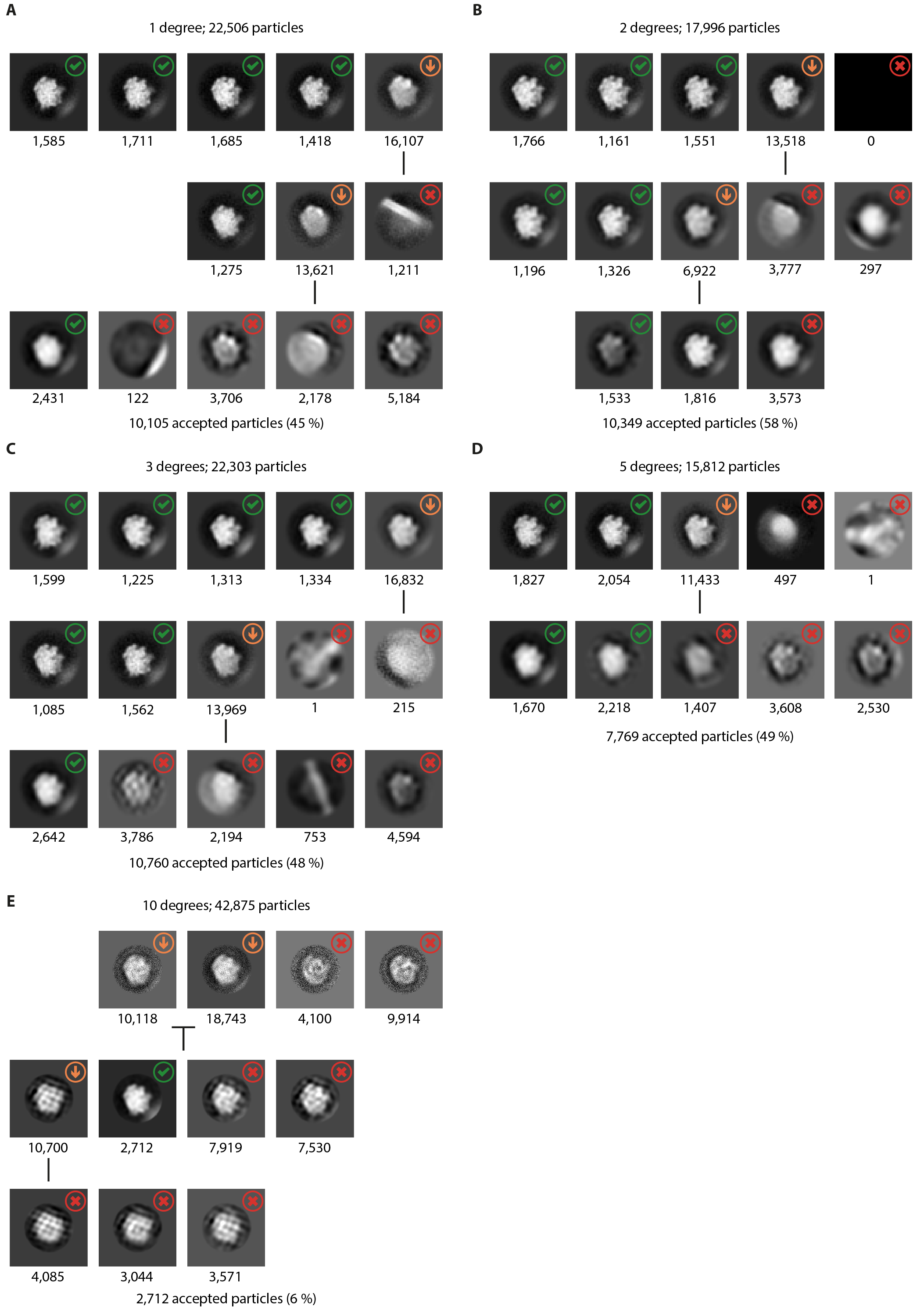

**SFig. 8 3D classification results after TM to remove junk particles.** Resulting classes after 3D classification to remove junk particles, performed on subtomograms with binning factor 6, corresponding to a voxel size of 11.8 Å/px, for tilt increments of 1 degree (**A**), 2 degrees (**B**), 3 degrees (**C**), 5 degrees (**D**), and 10 degrees (**E**). Z-score thresholds of 5 were used for all data, apart from the 10-degree data, where a threshold value of 3.75 was used. For the 10-degree dataset, a binning factor of 2 is used, corresponding to a voxel size of 3.9 Å. The total number of starting particles is indicated at the top. The number of particles per class is indicated below each class. Successive classification rounds are visible as rows. Particle classes that were discarded have been marked with a red arrow, particle classes that were taken to the next round of classification are indicated with a downward orange arrow, and particle classes that were accepted as true positives are indicated with a green check mark. The number of accepted particles after 3D classification is indicated below, as well as the percentage with respect to the number of starting particles.

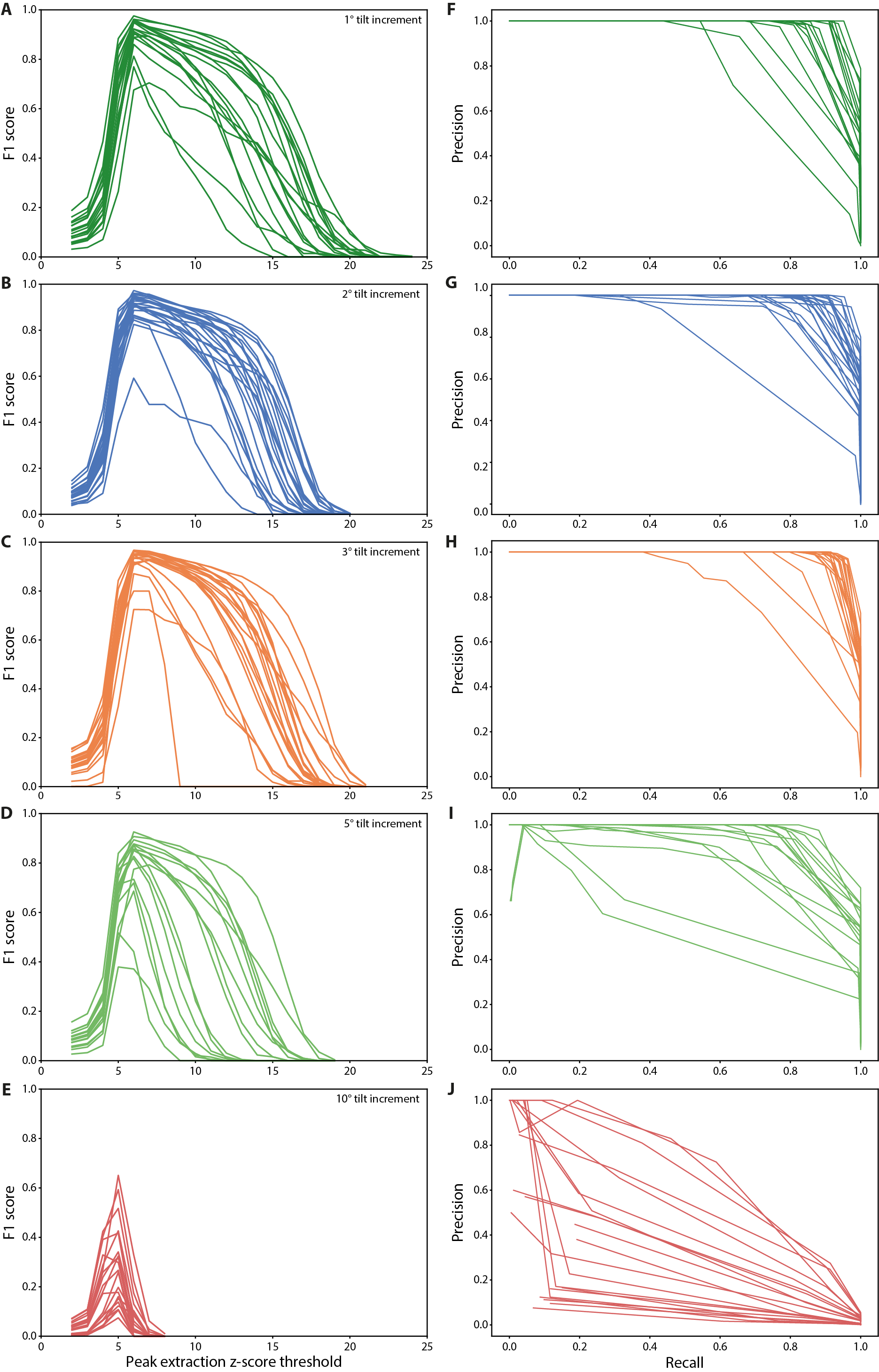

**SFig. 9 F1- and PR-curves for all tomograms**. Each line represents a single tomogram used in the analysis. **A-E** F1-curves for tomograms used for TM for 1-, 2-, 3-, 5-, and 10-degree increments respectively. On the horizontal axes the z-score threshold used for peak extraction is plotted and on the vertical axes the F1-score for particles extracted using that threshold. **F**-**J** Precision - recall curves for the same data.

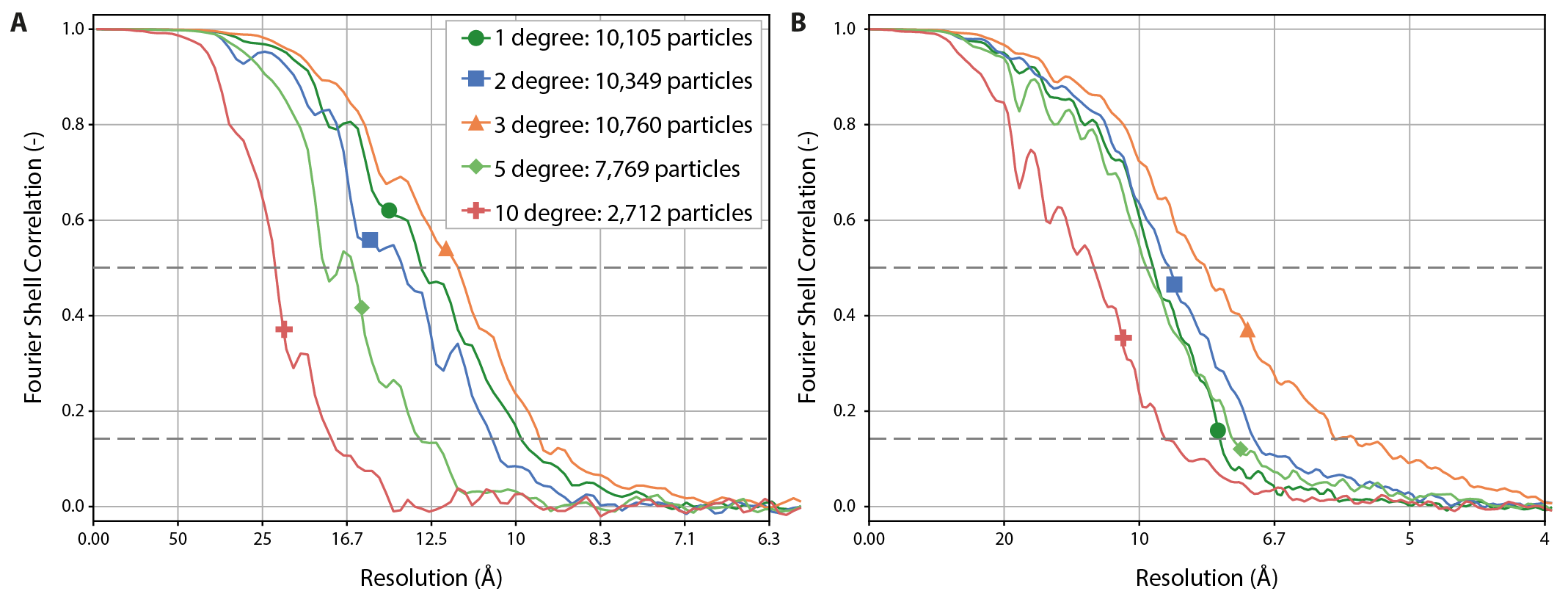

**SFig. 10 Fourier shell correlation curves**. **A** FSC curve for all tilt increments after refinement in RELION. **B** FSC curve for all tilt increments after refinement in M. Thresholds at FSC of 0.5 and 0.143 are indicated with dashed horizontal lines.

**Supplementary Note:**

As the tomogram SNR distribution of particularly the 2° and 3° dataset showed similar SNR distributions, we performed formal significance testing for the data in this panel (see Fig. 2B). Kruskal-Wallis test confirmed significant differences across conditions (H=270.97, p<0.001); pairwise Mann-Whitney U tests with Bonferroni correction revealed all pairs were significantly different except 2° vs. 3° (p=0.093), indicating comparable SNR for these two tilt increments.
